# Single mRNP analysis by super-resolution microscopy and fluorescence correlation spectroscopy reveals that small mRNP granules represent mRNA singletons

**DOI:** 10.1101/558098

**Authors:** Àngels Mateu-Regué, Jan Christiansen, Frederik Otzen Bagger, Christian Hellriegel, Finn Cilius Nielsen

## Abstract

Small cytoplasmic mRNP granules are implicated in mRNA transport, translational control and decay. Employing Super-resolution Microscopy and Fluorescence Correlation Spectroscopy, we analyzed the molecular composition and dynamics of single cytoplasmic YBX1_IMP1 mRNP granules in live cells. Granules appeared elongated and branched with patches of IMP1 and YBX1 distributed along mRNA, reflecting the attachment of the two RNA-binding proteins in *cis*. Particles form at the nuclear pore and are spatially segregated from translating ribosomes, so the mRNP is a repository for mRNAs awaiting translation. In agreement with the average number of mRNA-binding sites derived from CLIP analyses, individual mRNPs contain 5 to 15 molecules of YBX1 and IMP1 and a single poly(A) tail identified by PABPC1. Taken together, we conclude that small cytoplasmic mRNP granules are mRNA singletons, thus depicting the cellular transcriptome. Consequently, expression of functionally related mRNAs in RNA regulons is unlikely to result from coordinated assembly.

## Introduction

Essential steps from the nuclear mRNA processing and export to cytoplasmic localization, translation and decay of mRNAs are implicated in fine tuning of gene expression. Regulatory steps are governed by RNA-binding proteins, which interact with the mRNA in a sequential manner. Mammalian cells comprise about 1500 RNA-binding proteins of which nearly half bind to mRNAs (Gerstberger et al., 2014), and the majority is widely distributed with a minority exhibiting a particular spatial and temporal expression. Some mRNA-binding proteins shuttle between the nucleus and the cytoplasm, whereas others are mainly present in the cytoplasm (Shyu and Wilkinson, 2000), including proteins such as FMRP, HuD as well as YBX1 and the family of insulin-like growth factor 2 mRNA binding proteins IMP1, 2 and 3 (Darnell and Richter, 2012). Cytoplasmic mRNA-binding proteins are found in small membrane-less granules. In contrast to the larger P-bodies and stress granules, which mostly embody stress-induced condensations of RBPs and mRNA promoted via intrinsic disordered regions in the attached RBPs (Molliex et al., 2015; Reijns et al., 2008), small cytoplasmic mRNP granules represent the unperturbed state of cellular mRNA. Due to their significance in dendritic and axonal mRNA transport, cytoplasmic mRNP granules are sometimes referred to as neuronal RNP granules (Anderson and Kedersha, 2006; Kiebler and Bassell, 2006), but mRNP granules are found in any cell throughout the body at all developmental stages. Global biochemical analyses have shown that granules contain ribosomal subunits, translation factors, decay enzymes, helicases, scaffold proteins, and RNA-binding proteins, but we have limited data about the molecular composition of individual granules, that allow us to understand their function and relation to the other granular assemblies.

Insulin-like growth factor 2 (IGF2) mRNA-binding protein 1 (IMP1, IGF2BP1) belongs to a conserved family of heterochronic mRNA-binding proteins (IMP1, IMP2, and IMP3) (Hansen et al., 2004; Nielsen et al., 2001;Nielsen et al., 1999; Yaniv and Yisraeli, 2002). Together with the cytoplasmic mRNA-binding protein Y-box binding protein 1 (YBX1), IMPs form typical cytoplasmic RNP granules (Eliscovich et al., 2017) (Figure 1). Granules are mobile and widespread in the cytoplasm although they exhibit a preponderance for the perinuclear regions and the lamellipodia in motile cells (Nielsen et al., 1999; Oleynikov and Singer, 2003). In conventional laser scanning microscopy they exhibit an optical diameter of about 200-700 nm (Nielsen et al.,2002). Transcriptome-wide CLIP analyses have shown that both IMP1 and YBX1 associate with large parts of the transcriptome (Conway et al., 2016; Goodarzi et al., 2015). Individual RNAs exhibit numerous IMP1 and YBX1 attachment sites distributed along the target mRNA (Nielsen et al., 2004; Runge et al., 2000; Singh et al., 2015), although it is unknown whether binding is taking place on the same mRNA molecule. IMP1 and YBX1 are essential for normal development; IMP1- and YBX1-deficient mice are both small with imperfect organ development including neuronal defects (Hansen et al., 2004; Uchiumi et al., 2006), and at the cellular level both factors promote cell growth (Bell et al., 2013; Bommert et al., 2013; Shiota et al., 2008). Moreover, YBX1 and IMP1 have been implicated in a series of complex biological pathways such as F-actin formation and protein secretion (Jønson et al., 2007; Uchiumi et al., 2006), and specifically IMP1 is a participant in the embryonal heterochronic network consisting of *HMGA2, let-7* and *Lin28B* mRNAs (Jonson et al., 2014;Nishino et al., 2013). Finally, YBX1 plays a role in nodal signaling via *sqt* RNA localization, processing and translation (Kumari et al., 2013).

**Figure 1.**
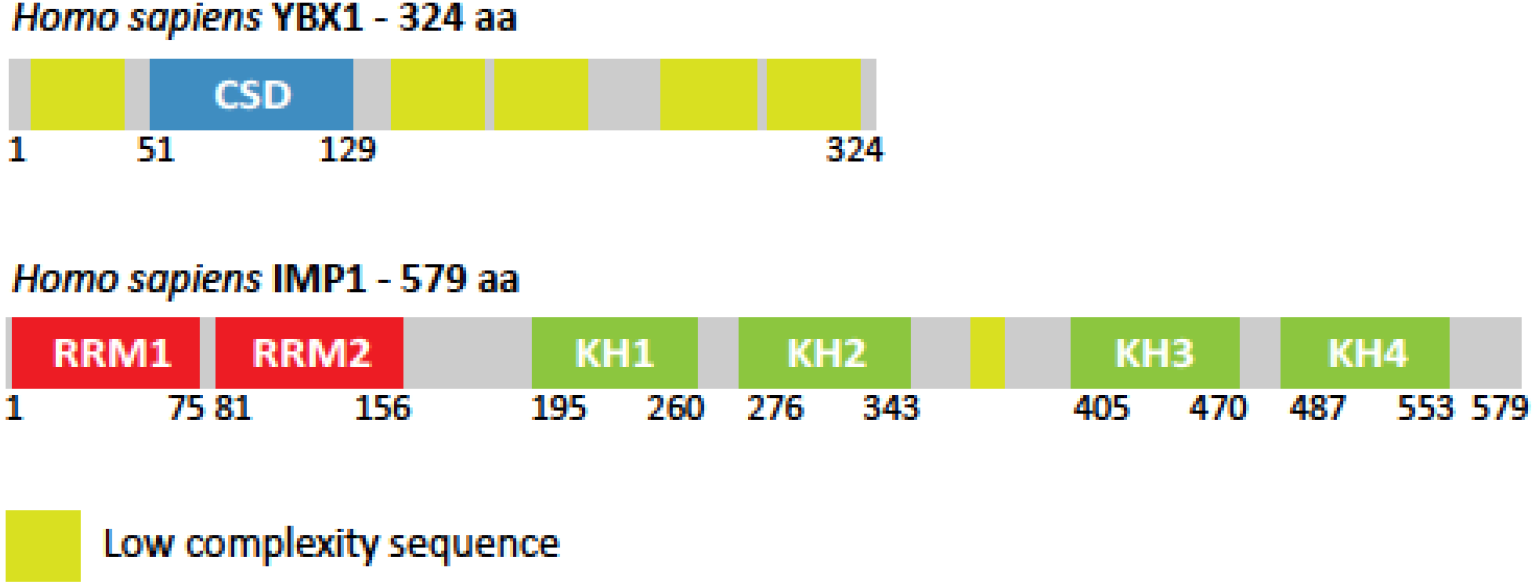
Schematic representation of YBX1 and IMP1 RNA-binding proteins. YBX1 is composed of a cold-shock domain (CSD) depicted in blue and a number of predicted low-complexity sequences (LCS) shown in yellow. IMP1 is composed of two RNA recognition modules (RRM) and four K homology domains (KH), the latter being responsible for IMP1 mRNA binding. Moreover, a short low-complexity sequence is predicted between domains KH2 and KH3.

To advance our understanding of cytoplasmic mRNP granules, we employed super-resolution and fluorescence correlation microscopy of YBX1, IMP1 and associated mRNA. In contrast to the current perception based on conventional light microscopy, our data show that granules represent irregular branched and elongated structures composed of alternating patches of IMP1 and YBX1 along a common mRNA. Formation of the particles requires mRNA, and mRNPs are first observed at the nuclear pore. The mRNPs are not connected to actively translating ribosomes, which are located in a separate vicinal compartment. Each particle contains a single mRNA and between 5-15 IMP1 and YBX1 molecules, in agreement with the average number of binding sites in the target mRNAs. Moreover, only a single poly(A) tail depicted by poly(A) binding protein C1 (PABPC1) staining was identified in each granule. Taken as a whole, we conclude that mRNP granules represent singletons and that coordinated expression of functionally related mRNAs is unlikely to be due to coordinated assembly.

## Results

### IMP1 and YBX1 coexist in mRNPs in an RNA-dependent manner

To provide an overview of the subcellular distribution and structure of IMP1 and YBX1 RNPs *in vivo*, HeLa cells were stained with anti-IMP1 and anti-YBX1 antibodies and examined by Structured Illumination Microscopy (SIM). IMP1 and YBX1 were cytoplasmic and prominent at the perinuclear region and in the lamellipodia of the cells (Figure 2A, panels 1-3). Granules appeared branched and elongated and ranged from 250 to 800 nm in size and were composed of alternating patches of IMP1 and YBX1 (Figure 2, panels 4-6). In general, YBX1 was observed along the entire outline of the particle, whereas IMP1 exhibited a preponderance for projections and ends. Consequently, single mRNP granules were defined by the coherent pattern of YBX1, which is also one of the most abundant protein components of mRNP granules (Singh et al., 2015). A comparison between the appearance of the mRNP granules in conventional Laser Scanning Microscopy (LSM) compared to SIM is shown in Supplemental Figure 6A. To visualize the associated mRNA, we performed FISH of *ACTB* mRNA and *GAPDH* mRNA combined with immunostaining of YBX1 and IMP1 (Figure 2B). *ACTB* mRNA and *GAPDH* mRNA were hybridized to 48 labelled Quasar^®^ 570 Dye-labeled probes covering the entire mRNA from the 5’ to the 3’ end. Probe staining was partially masked by the attached RNA-binding proteins, in particular when YBX1 was present. No granular overlap between *ACTB* and *GAPDH* mRNAs was observed. To corroborate the putative RNA-dependent interaction of YBX1 and IMP1, an immunoprecipitation of GFP-tagged IMP1 was carried out (Figure 2C). YBX1 was enriched in the immunoprecipitate, and treatment with RNAse A reduced the amount of immunoprecipitated YBX1, indicating that IMP1 does not interact directly with YBX1. In agreement with the observation that the particles contain multiple IMP1 molecules binding independently to the mRNA, endogenous IMP1 was also reduced by RNAse A (Figure 2C). The mRNPs were clearly distinct from cytoplasmic P-bodies and stress-granules identified by G3BP and DCP1a, respectively (Figure 2A, panels 7-8 and Supplemental Figure 1). The average size of stress-granules was at least an order of magnitude larger than IMP1_YBX1 mRNPs, that could be distinguished within the larger bodies.

**Figure 2.**
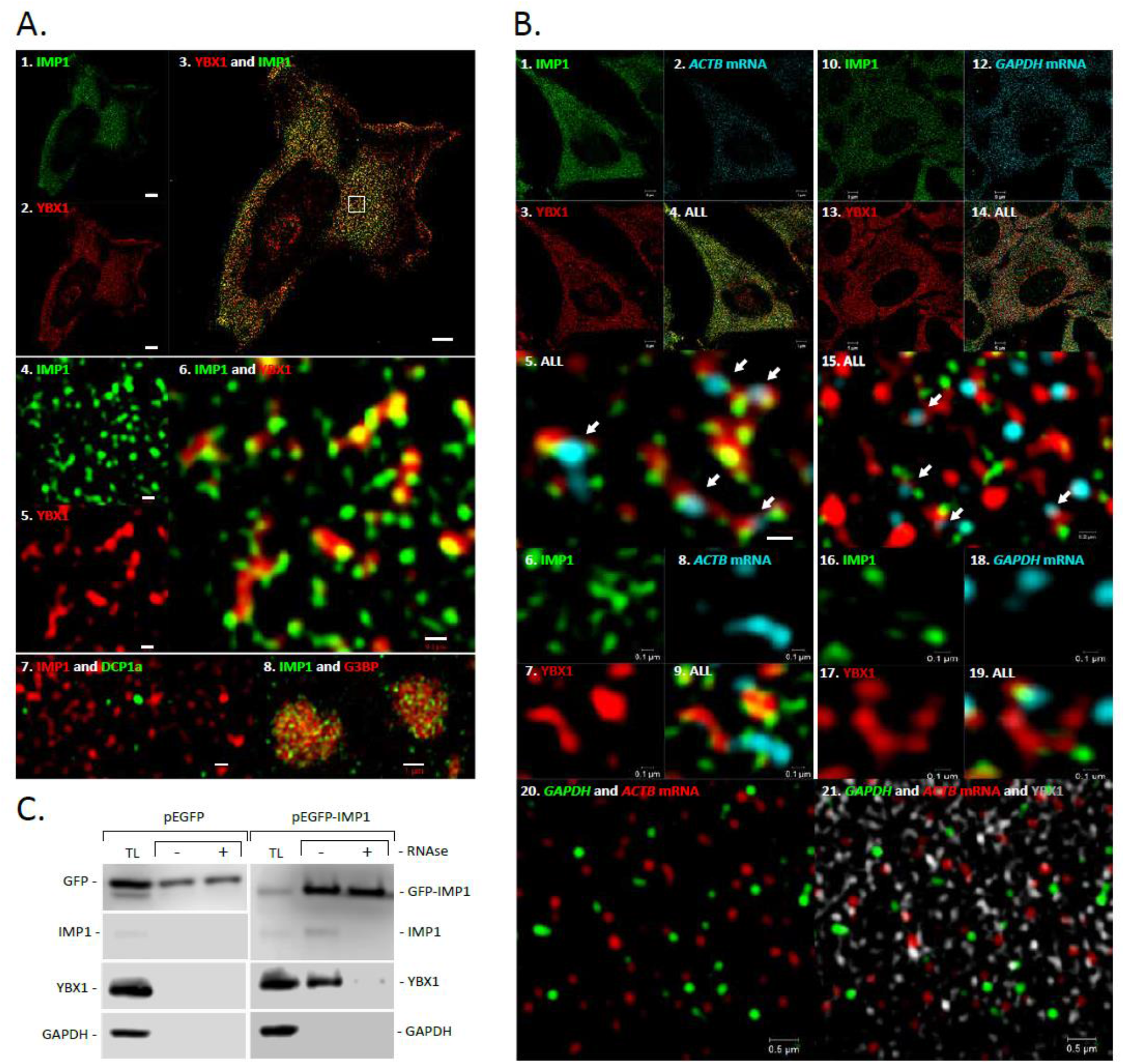
Structure of IMP1 and YBX1 and *ACTB* mRNA mRNP granules. **A1-6,** HeLa cells were stained with anti-IMP1 and anti-YBX1 antibodies followed by Alexa fluor 488 (green) and Alexa fluor 647 (red) secondary antibodies, respectively. **A1-3,** an overview of the cell. Scale bars = 5 μm. and **A4-6,** a blow up of IMP1 and YBX1 granules in the indicated area (white square) in panel **A3**. Scale bar = 0,2 μm. **A7,** P bodies depicted by DCP1a-EGFP in combination with IMP1 staining and **A8,** pHcRed-G3BP in stress granules in combination with IMP1 staining. **B1-9,** IMP1 (green, Alexa Fluor 488) and YBX1 (red, Alexa Fluor 647) immunostaining in combination with *ACTB mRNA* FISH (cyan) employing 48 Quasar-570 labelled oligonucleotides corresponding to the entire *ACTB* mRNA. **B1-4,** Overview of a HeLa cell. Scale bars = 5 μm. **B5-9,** blow up of *ACTB* mRNA and IMP1_YBX1 containing granules. Scale bars = 0,2 μm (**A5**) and 0,1 μm (**A6-9**). **B10-19,** IMP1 (green, Alexa Fluor 488) and YBX1 (red, Alexa Fluor 647) immunostaining in combination with *GAPDH mRNA* FISH (cyan) employing 48 Quasar-570 (cyan) labelled oligonucleotides corresponding to the entire *GAPDH* mRNA. **B10-14,** Overview of a HeLa cell. Scale bars = 5 μm. **B15-19,** blow up of *GAPDH* mRNA and IMP1_YBX1 containing granules. Scale bars = 0,2 μm (**B15**) and 0,1 μm (**B16-19**). **A20-21,** Double FISH with *ACTB mRNA* (red, Quasar-670 conjugated probes) and *GAPDH mRNA* (green, Quasar-570 conjugated probes) in combination with YBX1 immunostaining (grey, Alexa Fluor 488). **C,** EGFP immunoprecipitation of transiently transfected HeLa cells with pEGFP-C1 (control) and pEGFP-IMP1. Immunodetection of GFP and GFP-IMP1, endogenous IMP1, YBX1 and GADPH, respectively, was performed in total lysate and immunoprecipitated (IP) fractions without or with (-/+) RNAse A treatment.

### YBX1 and IMP1 mRNP is formed in the nuclear pore and awaits translation

Since the observed size of the mRNP is difficult to reconcile with facilitated diffusion of IMP1_YBX1 mRNP particles through the central nuclear pore channel, whose size limit is regarded to be about 40 nm (Hoelz et al., 2011), we examined the first appearance of the mRNP. SIM revealed very faint and almost non-existent YBX1 and IMP1 nuclear staining, and we failed to observe any colocalization between the two factors and mRNA. This led us to image the nuclear pore in closer detail. Nuclear pores were visualized by staining of Nup153, and both IMP1 and YBX1 mRNPs were found to align with the pore (Figure 3), where mRNPs projected towards the cytoplasm. The cytoplasmic distribution and alignment was unaffected by incubation with leptomycin (data not shown). Consequently, we infer that IMP1_YBX1 mRNPs are likely to form at the nuclear pore. To define the relation of the mRNP to the translation apparatus, we performed a polysome fractionation analysis, which showed that both IMP1 and YBX1 predominantly sedimented as free mRNP in monosomal fractions corresponding to 40S - 80S (Figure 4A). In addition, we employed O-propargyl-puromycin (OPP) to depict actively translating ribosomes (Liu et al., 2012). After fixation, OPP was conjugated to an Alexa Fluor-488 fluorophore by click chemistry before IMP1 and YBX1 were stained by immunofluorescence, and SIM Z-stacks of the cells were generated. As shown in Figure 4 (Panels B and C) and Supplemental Figure 3, IMP_YBX1 mRNP did not colocalize with translating ribosomes in agreement with the polysome analysis. However, ribosomes were positioned in close proximity to the mRNPs. Taken together, we the data indicate that mRNPS are not directly associated with translating ribosomes and that protein synthesis follows unloading of the mRNA.

**Figure 3.**
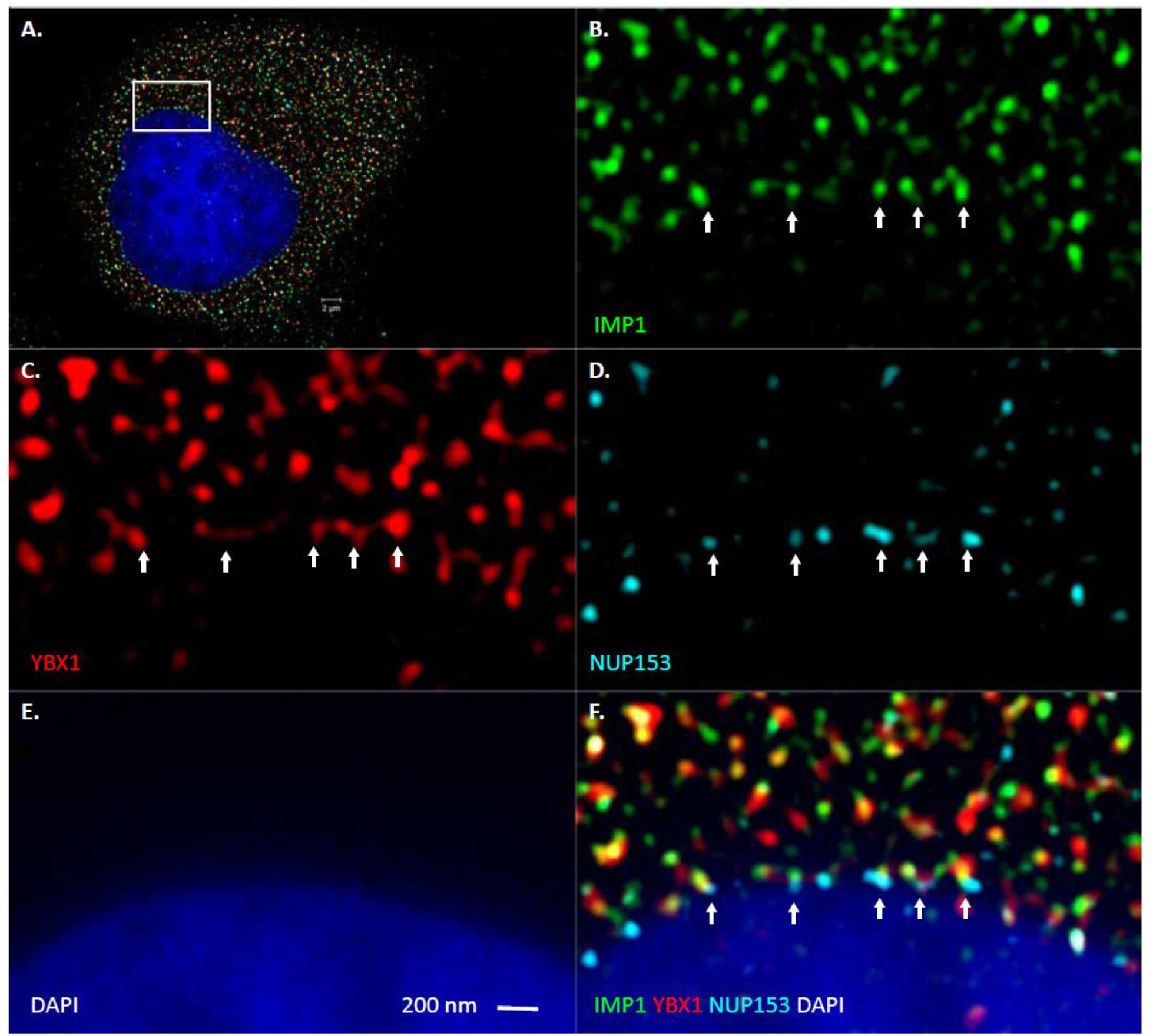
IMP1 and YBX1 mRNP formation at the nuclear pore. Hela cells were stained with anti-IMP1, anti-YBX1 and anti-NUP153 primary antibodies followed by Alexa fluor 488 (green), Alexa 555 (cyan) and Alexa fluor 647 (red) secondary antibodies, respectively. Moreover, the nucleus was stained with DAPI (deep blue). **A,** Overview of a triple-stained cell and the area that is shown in the blow up below. **BE,** Individual IMP1, YBX1, Nup153 and DAPI stainings. **F,** Composite picture demonstrating the colocalization of Nup153 and the IMP1_YBX1 mRNP. Arrows indicate nuclear pores.

**Figure 4.**
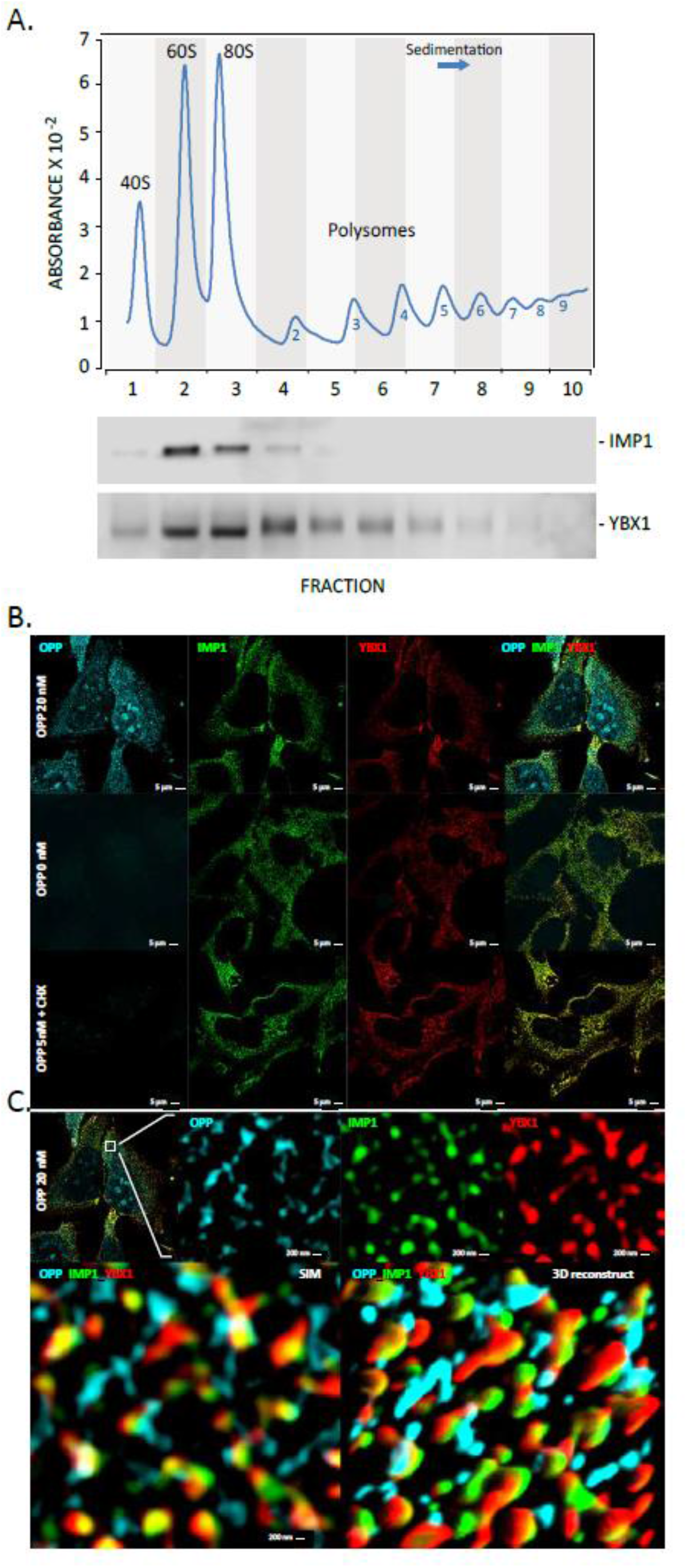
IMP1 and YBX1 mRNP granules and the translational apparatus. **A,** Polysome fractionation analysis and subsequent immunodetection of IMP1 and YBX1 by Western Blot. The upper panel depicts the A260 sedimentation profile, whereas the lower panel shows the corresponding Western analysis of the fractions. The sedimentation of 80S monosomes and ribosomal subunits is indicated, and the polysomes are numbered according to the number of loaded 80S complexes. **B,** Translating ribosomes were labelled with O-propargyl puromycin (OPP) followed by puromycin-based click conjugation of Alexa Fluor 488 (cyan). IMP1 and YBX1 were stained with anti-IMP1 and anti-YBX1 antibodies followed by Alexa Fluor 568 (green) and Alexa Fluor 647 (red) secondary antibodies, respectively. **A,** OPP, IMP1 and YBX1 stainings with and without addition of 20 nM OPP and in the presence of cycloheximide (CHX). **C,** Blow up of the individual mRNP and 3D reconstruction of 10 consecutive slices demonstrating the vicinity of ribosomes and mRNP.

### Dynamics of IMP1_YBX1 mRNPs

To assess the dynamics of cytoplasmic IMP1_YBX1 mRNPs, we employed Fluorescence Correlation Spectrosopy (FCS), which allows quantification of diffusion time, concentration and brightness of molecules (or particles) in solution or in live cells. Wild-type GFP-tagged IMP1 and -YBX1 and a GFP-tagged IMP1 GXXG mutant (GFP-IMP1_KH1-4mut), with impaired RNA-binding (Supplemental Figure 4), were expressed in HeLa cells. Moreover, GFP was included as a reference. As depicted in Figure 5A, GFP and GFP-IMP1_KH1-4mut exhibited faster Lag times than GFP-IMP1 and GFP-YBX1. Whereas GFP best fitted a 1-component model, the best fit of the experimental autocorrelation curves of GFP-IMP1_KH1-4mut, GFP-IMP1 and GFP-YBX1 was a 2-component model. The modelling showed that GFP-IMP1_KH1-4mut could be resolved into two relatively fast diffusion times of 2.9E-04 s and 1.2E-03 s, respectively, whereas GFP-IMP1 and GFP-YBX1 exhibited a fast moving fraction with a Lag time of ~10E-03 s and a much slower fraction moving in the range of 10E-01 s (Figure 5C). Compared to GFP, wild-type IMP1 and YBX1 exhibited fractions diffusing 10 fold and a 1000 fold slower. In agreement with the pull-down analysis described above, Fluorescence Cross Correlation Spectroscopy with GFP-IMP1 or GFP-IMP1_KH1-4mut combined with mCherry-YBX1 demonstrated that the interaction between the two factors *in vivo* relies on mRNA-binding and not on other factors (Figure 5B). To further substantiate FCS data, we visualized the mobility of the RNP in the live cell. We employed mEos3.2, a photoconvertible fluorescent protein (Zhang et al., 2012), and expressed mEos3.2-IMP1. After localized photoactivation at 405 nm, cellular movements of the photoconverted red mEos3.2-IMP1 was followed by time lapse microscopy (Supplementary Figure 4). Both a slow and a rapid transport (in the range of seconds) was observed. Slow particles diffused in all directions and migrated at very slow rate of about 0.1 μm/s. Even after 1.5 minute the majority of the granular RNP remained in the center of the photoactivation. In contrast, the rapid diffusion accumulated quickly at a particular distant site 25 μm away within seconds after photoconversion, corresponding to a speed of at least ~5 μm/s. The results show that assembly of granules requires RNA-binding, and that IMP1_YBX1 mRNP motility is multidirectional and involves fast and slow trafficking and regulated anchoring.

**Figure 5.**
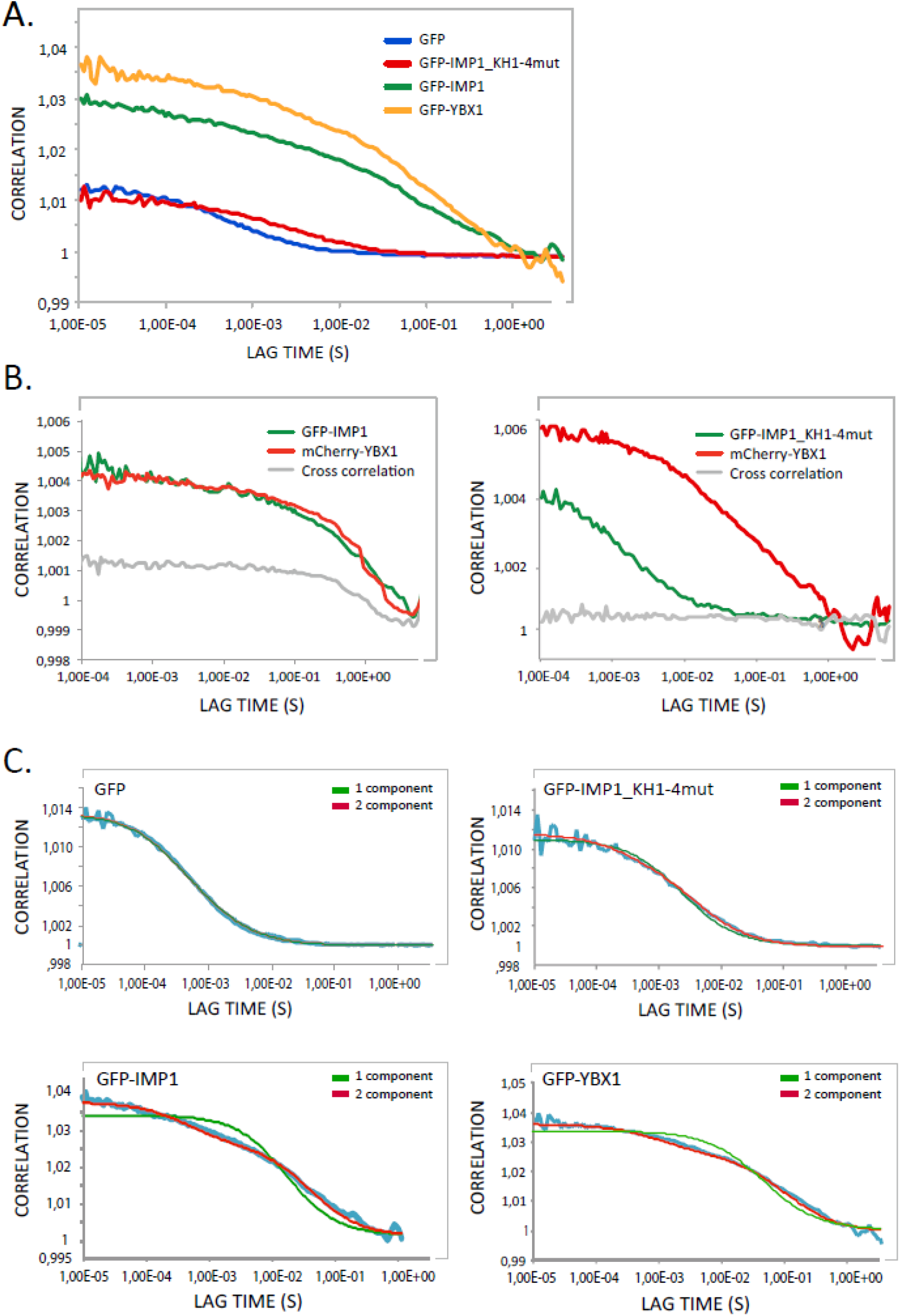
IMP1_YBX1 mRNP dynamics. GFP- and mCherry-tagged IMP1, YBX1 and IMP1 mutated in the four GXXG loops (GFP-IMP1_KH1-4mut) were expressed in HeLa cells, and their dynamic behaviour was recorded by Fluorescence Correlation Spectroscopy (FCS). **A,** Autocorrelation curves of cytoplasmic GFP, GFP-IMP1, GFP-YBX1 and GFP-IMP1_KH1-4mut. **B,** Cross-Correlation curves of cells co-transfected with GFP-IMP1 and mCherry-YBX1 or co-transfected with GFP-IMP1_KH1-4mut and mCherry-YBX1. **C,** Autocorrelation curves of the different constructs shown in panel A together with the fittings to 1- and 2-component diffusion models.

### Molecular composition of IMP1_YBX1 mRNP

We analysed the number of molecules in IMP1_YBX1 mRNP *in vivo* by means of Fluorescence Correlation Spectroscopy (FCS) and Localization Microscopy and compared the results to available transcriptome-wide IMP1 and YBX1 eCLIP and RIP-seq data. Moreover, the interplay between IMP1 and YBX1 upon binding to high-affinity RNA targets such as *ACTB* and *C-MYC* mRNAs (Leeds et al., 1997; Ross et al., 1997) was examined in order to provide a framework for an understanding of the stoichiometric data (Figure 6).

**Figure 6.**
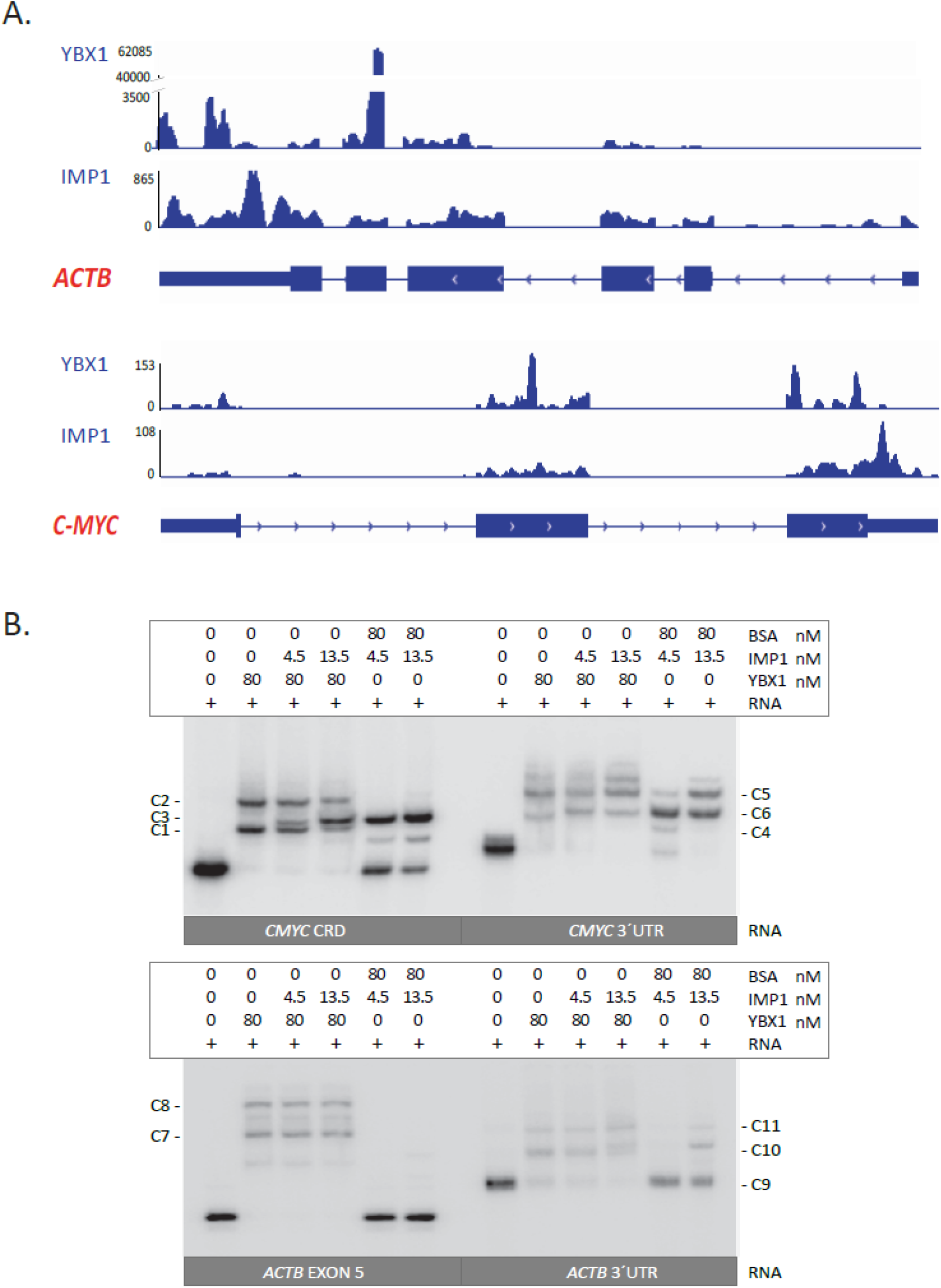
RNA-binding properties of IMP1 and YBX1. **A,** YBX1 RIP-seq and IMP1 eCLIP data from *ACTB* and *C-MYC* mRNAs. **B**, Electrophoretic mobility-shift assay (EMSA) of single or combinations of IMP1 and YBX1 proteins with ^32^P-labelled *C-MYC* coding region determinant (CRD), *C-MYC* 3’UTR, *ACTB* Exon5, or *ACTB* 3’UTR.

Figure 6A shows a schematic representation of *ACTB* and *C-MYC* transcripts together with the CLIP-derived number of cross-linked and immunoprecipitated reads at particular positions (Conway et al., 2016; Goodarzi et al., 2015). Although IMP1 exhibits a preponderance for the 3’UTR and YBX1 binding is more widespread, some binding sites are overlapping and the two factors may compete for a number of binding sites. The CLIP data show that YBX1 and IMP1 exhibit strong binding to *ACTB* mRNA exon 5 and the ZIP code in the 3’UTR, respectively, and to the CRD region and the 3’UTR of *C-MYC*, respectively. Since CLIP experiments provide a global overview of binding sites and are unable to distinguish between *cis*- and *trans*-attachments, we supplemented the CLIP data with electrophoretic mobility-shift assays (EMSA) with these mRNAs, shown in Figure 6B. Albeit *in vitro*, the advantage of EMSA is the ability to examine putative *cis*-attachments by identifying supershifts, and the chosen YBX1 and IMP1 concentration ratios were according to the protein copy number per HeLa cell, as previously characterised (Singh et al., 2015). YBX1, IMP1 or both proteins were incubated with the radioactively labelled RNA targets. The mobility-shifts showed that the binding of each protein to the chosen mRNA segments was mutually exclusive, since there was no evidence of a supershift with any of the RNA probes. For *C-MYC*, we observed that both YBX1 and IMP1 were able to bind to the CRD and 3’UTR targets. IMP1 exhibited a higher affinity for CRD than YBX1, since at 1:10 of the YBX1 concentration IMP1 was able to out-compete YBX1. The same happened with *C-MYC* 3’UTR, although in this case IMP1 was able to achieve a higher degree of multimerization. Regarding the *ACTB* segments, we observed that YBX1 had a very high affinity for exon 5 and showed a high degree of multimerization, in accordance with the CLIP data. IMP1 was essentially unable to bind to *ACTB* exon 5 and could not compete with YBX1. Both IMP1 and YBX1 were able to bind to the 3’UTR target, and we observed the same competition pattern as described for the *C-MYC* transcripts, although both proteins exhibited lower affinity. Taken together, we infer that IMP1 and YBX1 compete for shared binding segments – so at a given time IMP1 or YBX1 may not occupy all their putative binding sites.

To determine the number of IMP1, YBX1 and RNA molecules in the individual mRNP, we employed Localization Microscopy and Fluorescence Correlation Spectroscopy. For Localization Microscopy, HeLa cells were stained with anti-IMP1 or anti-YBX1 antibodies and Alexa Fluor 568 or 555 secondary antibodies, respectively, or with a set of *ACTB* mRNA probes as shown in Figure 2. The number of secondary antibodies binding a primary antibody was determined by FCS, and this showed that two secondary antibodies bind to each primary antibody (data not shown). Consequently, events (emitted photons at a particular site) were divided by two (Figure 7C). To avoid counting the same secondary antibody more than once due to the presence of multiple fluorophores and to improve positioning accuracy, a grouping of 5.5 pixels was applied. Panel A shows Localization Microscopy readings with GAUSS distributions and crosses. Clustered photons (crosses) corresponding to 50 granules were counted (Figure 7A), and the median and range are summarized in the Table (Figure 7C). The median number of molecules was 7 for YBX1 (range 2-34) and 6 for IMP1 (range 2-15) in the RNP. Moreover, we analyzed the number of *ACTB* RNA probes in the mRNP granules. As illustrated in Figure 2, parts of the *ACTB* mRNA appeared to be masked by attached RBPs, and we counted on average 11 probes ranging from 3-42 in the mRNP. In no *ACTB* cluster, did we count more than the maximal number of 48 probes indicating that there is only a single *ACTB* transcript in a particle.

**Figure 7.**
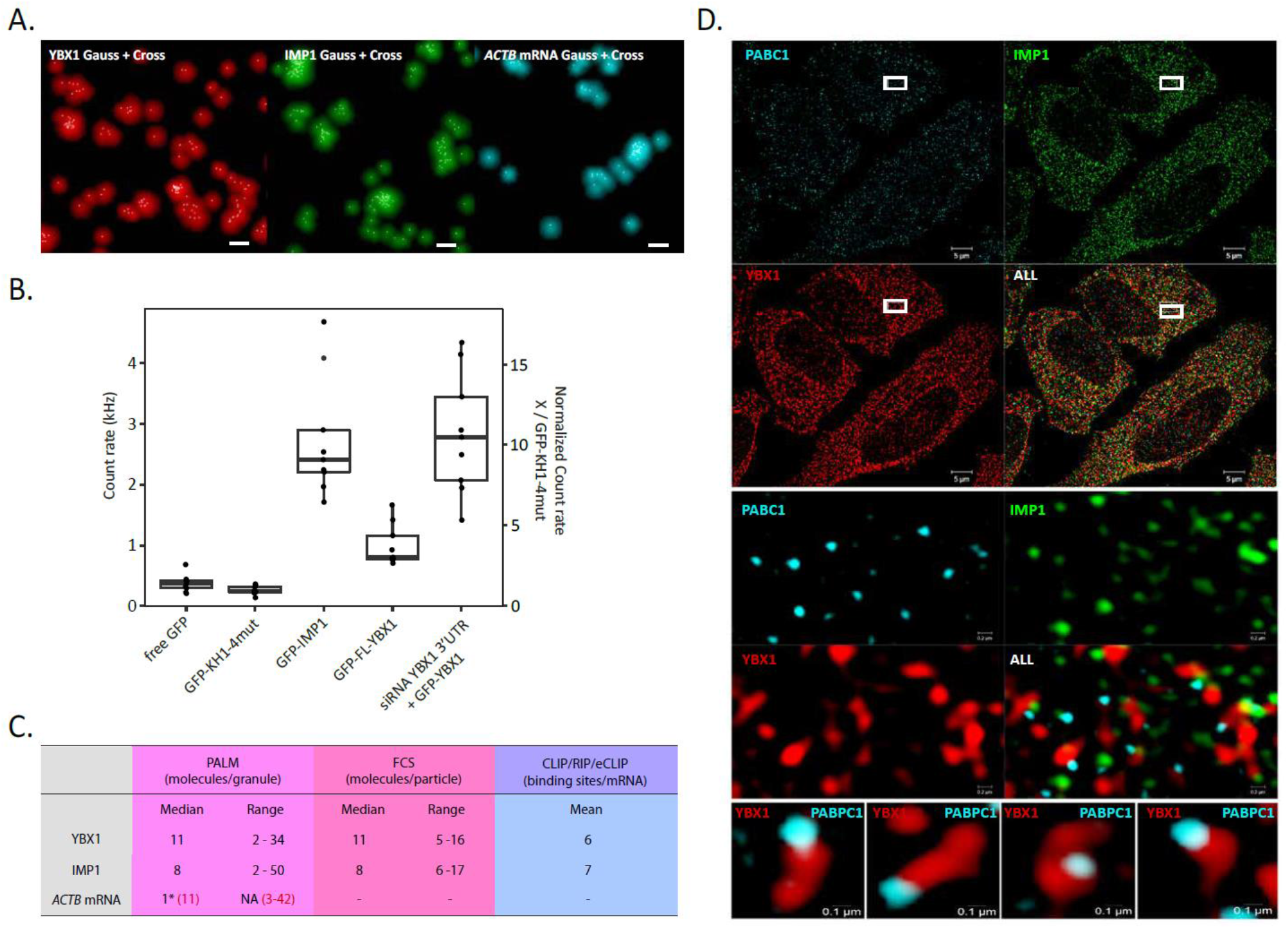
Molecular composition of IMP1_YBX1 mRNP. The number of IMP1, YBX1 and *ACTB* mRNA molecules in the mRNP were derived from Localization Microscopy (LM) or from FCS and compared to the average number of binding sites in the transcriptome from eCLIP and RIP-seq analysis. For Localization Microscopy cells were stained for IMP1, YBX1 and *ACTB* mRNA, and following bleaching mRNP emitted photons were counted as described. **A**, Examples of the Localization Microscopy images of YBX1 (red), IMP1 (green) and *ACTB* mRNA (cyan), respectively. Scale Bar = 100 nm **B,** Counts per particle derived from Fluorescence Correlation Spectroscopy of cells transfected with GFP, GFP-IMP1_KH1-4mut, GFP-IMP1, GFP-YBX1, and with co-transfection of YBX1 3’UTR-directed siRNA and GFP-YBX1. Laser power was 0.02% in all measurements. **C**, Summary of the data from Localization Microscopy, FCS, and binding sites predicted from either eCLIP (IMP1) or RIP-seq (YBX1) experiments. **D,** Immunofluorescence staining of PABPC1 (cyan, Alexa Fluor 488), IMP1 (green, Alexa Fluor 568) and YBX1 (red, Alexa Fluor 647). An overview of a whole HeLa cell is shown in panel above (scale bar = 5 μm) followed by a blow-up image of the triple PABPC1, IMP1 and YBX1 staining of the area squared in the panel above is shown below (scale bar = 0,2 μm). Positioning of PABPC1 (cyan) in individual mRNPs depicted by YBX1 (red) staining are shown in the panel below (scale bar = 0,1 μm).

To corroborate the findings described above, we extracted information regarding the average number of molecules in the particles in *live* cells by FCS. Total count rate (kHz), correlation and counts per molecule data were obtained for each measurement. Total count rate refers to the total number of photons per second collected by a detector and correlation relates to the amplitude of the autocorrelation curve and was used to estimate concentration. The parameter referred as counts per molecule (or particle, kHz) is defined as the total count rate (kHz) divided by the number of molecules (or particles) determined by the amplitude of the autocorrelation curve. Counts per particle from free GFP and GFP-IMP1_KH1-4mut (monomers) were compared with the wild type GFP-IMP1 and GFP-YBX1 in HeLa cells. We could measure differences in the fluctuations during the recorded measurements (Figure 7B), and this was also represented in the amplitude of the autocorrelation curves. Cytoplasmic readings from 9 different cells showed that the number of GFP-IMP1 molecules per particle was from 7 to 11, considering GFP-IMP1_KH1-4mut as the monomer state of the protein, whereas the results for GFP-YBX1 were unexpectedly low compared to what we observed in the Localization Microscopy analysis. Endogenous YBX1 concentration is 10 times higher than IMP1, and we suspected that GFP-YBX1 counts per particle were underestimated due to the high levels of competing endogenous protein. To examine if this was the case, endogenous YBX1 was knocked down using a siRNA against the 3’UTR. As observed in Figure 7, counts per particle during these conditions increased from 3-4 to 8-13 YBX1 molecules per particle.

Finally, we estimated the number of global IMP1 and YBX1 mRNA binding sites from public CLIP and RIP-seq data (Conway et al., 2016; Goodarzi et al., 2015) in order to put the Localization Microscopy and FCS data in perspective (Figure 7, Panel C). Raw CLIP-seq data (fastq) were downloaded from the short read archive (SRA) and aligned to the human genome sequence. Binding sites were estimated by read-islands. The lower detection level was adjusted to the nearest threshold, where islands in the mock control samples equalled the total number of transcripts, therefore a cut-off of 7 and 6 for IMP1 and YBX1, respectively, was employed. We estimated that the average number of IMP1 and YBX1 binding sites per mRNA to be 7 and 5, respectively. IMP1 has been described to dimerize to form stable RNA-binding complexes (Nielsen et al., 2004), corroborated by the electrophoretic mobility-shift analysis in Figure 6, and YBX1 has also been shown to multimerize on a single RNA attachment site (Figure 6B) (Skabkin et al., 2004). Taken as a whole, Localization Microscopy and FCS analyses, which provide the molecular composition of the individual RNP granules *in vivo*, are in agreement with the transcriptome-wide CLIP estimates, when dimerization of IMP1 and a multimerization of YBX1 on mRNA are taken into account.

To further substantiate that the particles only contained a single mRNA, we employed a generic and global approach and determined the number of poly(A) tails in the particles. This was done by staining PABPC1, which represents the cytoplasmic poly(A) binding protein, in combination with YBX1 to depict the particles (Figure 8D). In no case did we observe more than one focus of PABPC1 per mRNP. Moreover, in the majority of the particles, the PABPC1 staining was characteristically located at the end of the particles. The observation was further substantiated by FCS measurements of PABPC1 showing that each PABPC1 foci consisted of 3-4 molecules of PABPC1 (Supplemental Figure 5) in agreement with the length of poly(A) tails in HeLa cells, which has been determined to contain from 50 to 100 nt (Chang et al., 2014), and taking into consideration that PABPC1 binds to a poly(A) tail every 27 nt (Baer and Kornberg, 1983). The diffusion rate of PABPC1-GFP was identical to that of YBX1 and IMP1 and cross-correlation was observed between the factors, indicating an *in vivo* interaction of the proteins (Supplemental Figure 5). We therefore conclude that IMP1_YBX1 RNPs represent solitary mRNAs with attached proteins.

## Discussion

Cytoplasmic RNP granules have been recognized for several decades, and although we have a relatively deep understanding of their biochemistry, the molecular composition of individual mRNPs is incompletely understood. This has partly been due to the lack of technologies to characterize molecular complexes in intact and live cells, so we employed super resolution microscopy and correlation spectroscopy to provide a deeper insight into the nature of single mRNP granules.

Compared to conventional laser scanning microscopy, where mRNP granules appear spherical with a size of 200-700 nm (Nielsen et al., 2002) (Supplemental Figure 6), structured illumination microscopy (SIM) has a resolution 2-3 times below that of diffraction-limited instruments, providing a lateral resolution of about 100 nm (Stelzer, 2014). An average 2 kb mRNA has an outline of about 300 nm - taking secondary structure into account - so it is feasible to distinguish RBPs and their position on the same mRNA by SIM (Milo et al., 2010) (http://book.bionumbers.org/which-is-bigger-mrna-or-the-protein-it-codes-for/), since the size of the complex is larger than the resolution. IMP1_YBX1 mRNPs were on average 300 nm in size, which is in agreement with atomic force microscopy of isolated granules (Jønson et al., 2007). YBX1- and IMP1-stainings were partly overlapping but also alternating along the mRNP. Moreover, the SIM analysis was supplemented by EMSA with four RNA targets of about 200 nucleotides, showing mutually exclusive attachment of YBX1 and IMP1, implying that the overlapping stainings observed in SIM are due to the resolution limit of about 100 nm. Whereas YBX1 was distributed along the entire particle, IMP1 had a preponderance for projections and ends. YBX1 is one of the core proteins of mRNPs and has previously been described to coat the entire mRNA (Singh et al., 2015; Skabkin et al., 2004), whereas IMP1 preferentially binds to single-stranded CA-rich elements in the 3’UTR and loop regions (Conway et al., 2016; Hafner et al., 2010). As previously described, the embedded mRNAs were to a large extent masked by the associated proteins (Buxbaum et al., 2014). In our data, mRNA appeared to be masked particularly in regions covered by YBX1, which is in accordance with the fact that YBX1 acts as a translational repressor (Evdokimova et al., 2006). Due to the distinct localization of the core proteins in the mRNP, the complete structure was only perceived in the composite pictures. If focus had been directed towards one of the factors, we would have failed to recognize the entire size and shape of the mRNP. Moreover, SIM directly demonstrates that YBX1 and IMP1 bind in *cis* like pearls on a string, which is of significance for the interpretation of CLIP and immunoprecipitation analysis that fail to make a distinction between *cis* and *trans*. The pattern of IMP1 and YBX1 in the particle is also in agreement with the mutually exclusive binding of the factors demonstrated in the band-shift analysis.

RNA-binding proteins associate with and dissociate from the mRNA along its journey from the nucleus to the translating ribosomes (Singh et al., 2015). Based on the presence of nuclear export signals in IMP1 and YBX1, the general idea has been that the factors enter the nucleus and bind their target mRNAs (Jung et al., 2018;Nielsen et al., 2003; Oleynikov and Singer, 2003). We hardly observed nuclear IMP1 and YBX1 staining, and proper mRNP complexes composed of IMP1 and YBX1 and mRNA were only identified at the cytoplasmic side of the nuclear pore. This observation supports a recent study, indicating that *Actb* mRNA first associates with IMP1 in the nuclear envelope (Wu et al., 2015). The nuclear pore is flexible and dynamic (Knockenhauer and Schwartz, 2016), and the largest macromolecular complexes that have been shown to pass are viral capsids up to ~40 nm diameter. Our data reflect that the mRNA is brought to the pore by a canonical mRNA export pathway, before IMP1 and YBX1 are loaded onto the mRNA. In this way, the nuclear pore may represent a crucial remodeling step of the mRNP (Singh et al., 2015). Whether the nuclear export signals in IMP1 and YBX1 have a function in localization to the nuclear pore remains to be addressed.

In the cytoplasm, IMP1 protects mRNAs from miRNA-mediated degradation (Jonson et al., 2014) and YBX1 is also an mRNA stabilizer (Evdokimova et al., 2001), so the mRNP is considered an mRNA repository/safe house until translation. Although granules incorporate ribosomal subunits (Krichevsky and Kosik, 2001), we observed no direct association with translating ribosomes, which were closely intertwined between the mRNPs. Our earlier compositional analysis of IMP1 RNPs, that identified the exon-junction components together with the nuclear cap-binding subunit CBP80 and the nuclear poly(A)-binding protein PABPN1, corroborates the pre-translational status of these cytoplasmic mRNPs (Jønson et al., 2007). In agreement with their large composite nature, the majority of the mRNPs exhibited a very slow diffusion rate compared to the KH domain GXXG loop mutant that failed to bind RNA. Particles migrated in all directions, and judged from the photoconversion experiments some were almost immobile, whereas about one-third of the mRNPs exhibited a faster mobility. The fastest particles were able to migrate several micrometers per second. Intriguingly, they accumulated at a particular location indicating that they may be subject to a regulated cytoplasmic docking. Transport may involve simple diffusion but also cytoplasmic streaming and motors, which are not mutually exclusive mechanisms (Lu et al., 2016; Song et al., 2015; Suzuki et al., 2017). Moreover, the single photoconverted mRNP was fairly stable, indicating that fluidity is low in contrast to what is observed within liquid droplets (Courchaine et al., 2016).

We determined the molecular composition of the individual granules by Localization Microscopy and FCS. Both methods roughly led to the same result and showed that particles were composed of 5-15 molecules of both IMP1 and YBX1. Due to the masking, the Localization Microscopy analysis of *ACTB* mRNA should obviously be interpreted with some caution. However, since we never arrived at probe counts exceeding the total number of applied *ACTB* probes, it suggested that there was only one mRNA in the particles and this was also reinforced by the observed lack of colocalization of abundant *ACTB* and *GAPDH* mRNAs in the granules. At a more global level, the observations were finally corroborated by the demonstration of only one PABPC1 staining focus in the distal parts of the granules. FCS data showed that each particle contained 3 to 4 PABPC1 molecules, in agreement with previous observation showing that the bulk of poly(A) tails in HeLa cells are 50-100 nt long (Chang et al., 2014) and that a single PABPC1 occupies 20 nt (Baer and Kornberg,1983). Finally, the comparative data from CLIP analysis (Conway et al., 2016; Goodarzi et al., 2015) show - in agreement with the Localization Microscopy and FCS data - that the average number of YBX1 and IMP1 mRNA attachment sites at a global level are 6 and 7, respectively. The reason that FCS and Localization Microscopy provide slightly higher numbers of IMP1 molecules is due to IMPs binding as dimers and that YBX1 to some extent multimerizes. The results are in line with findings showing that neuronal mRNAs travel singly into dendrites (Batish et al., 2012) and that *MAP2, CaMKIIa* and *ACTB* RNAs localize independently in low copy numbers (Mikl et al., 2011), thus supporting the “sushi-belt model” (Doyle and Kiebler, 2011). The findings may also be supported by a rough estimate. Assuming that a HeLa cell contains about 200.000 RNA polymerase II transcripts in a volume of 3000 μm^3^, then there will be 60 mRNAs per μm^3^(Shapiro et al., 2013). By simple counting of SIM stacks, we observe 30-50 mRNPs per μm^3^, which is compatible with a single transcript in each mRNP, considering that some mRNAs may undergo translation or reside in the nucleus.

RNA-binding proteins have been proposed to coordinate the production of functionally related proteins by organizing their mRNAs in regulons (Keene, 2007). The finding that small cytoplasmic mRNP granules represent singletons implies that coordinate expression of functionally related mRNAs in RNA regulons is unlikely to result from coordinated assembly, but rather results from regulated docking, as described above, or from selective stabilization of mRNA in the particles. Small cytoplasmic mRNP granules are distinct from large stress granules and P bodies, that represent dynamic mRNP assemblies formed in response to stress and mRNA decay, respectively. Based on our findings, we propose that smaller cytoplasmic granules should be designated mRNP singletons rather than granules to clearly distinguish them from the larger assemblies. Moreover, this would allude to the fact that stress granules probably incorporate elements of mRNP singletons.

## Acknowledgements

The Danish National Program for Infrastructure is thanked for donating the super resolution microscope and Lena Bjørn Johansson is thanked for technical assistance.

## Author Contributions

FCN and AM designed the study. AM, FCN, JC, FOB, CH designed the experiments. AM, FOB and JC performed the experiments. AM, FCN, JC, FOB and CH analysed the results. AM, FCN and JC prepared the manuscript. All authors read and approved the final manuscript.

## Methods

### Contact for Reagent and Resource Sharing

Further information and requests for resources and reagents should be directed to and will be fulfilled by the Lead Contact, Finn Cilius Nielsen.

### Experimental model and Subject Details

#### Cell lines

HeLa cells (ATCC^®^ CCL-2^™^) were cultured in phenol red-free Dulbecco’s Modified Eagle Medium (DMEM), high glucose (4.5 g/L) + GlutaMAX and 1 mM sodium pyruvate (Thermo Fisher Scientific) supplemented with 10% Fetal bovine serum (Tetracycline free, Biowest) and penicillin/streptomycin (Invitrogen). Cells were grown at 37°C with 5% CO_2_ in a humidified incubator.

### Method Details

#### Transfections

##### Vectors

IMP1 was cloned into pEGFP-C2 (Clontech) and pmEos3.2-C1 vector (Addgene). GFP-IMP1_KH1-4mut construct was obtained by mutating the GXXG loops of the 4 KH domains (Hollingworth et al., 2012) from GK(E/K/G)G to GELG. YBX1 was cloned into pcDNA3.1+N-EGFP (Genscript) and pmCherry-C1 (Clontech) vectors, inserting a 25 amino acid flexible linker composed of 5x GlyGlyGlyGlySer between the fluorescent tag and YBX1 to reduce aggregation. PABPC1 was cloned into pEGFP-N1, thus placing the GFP tag in the C-terminal of the protein. DCP1a and G3BP were cloned into pcDNA3-EGFP and pHcRed1-c1, respectively.

#### Plasmid transfections

HeLa cells were plated in 35mm glass-bottom dishes (P35G-1.5-14-C, MatTek) and were transfected using FuGENE^®^ 6 (Promega). For each transfection, 3 μl of FuGENE^®^ 6 and 1.3 μg of plasmid were added to 100 μl of OPTI-MEM Medium (Thermo Fisher Scientific). The mixture was incubated at room temperature for 30 minutes prior to the addition to the coverslips.

#### siRNA and plasmid co-transfection

HeLa cells were plated in 35mm glass-bottom dishes (P35G-1.5-14-C, MatTek) and were transfected using Lipofectamine 2000 (Thermo Fisher Scientific). For each transfection, 3 μl of Lipofectamine^®^ 2000, 1 μg of plasmid and siRNA to a final concentration of 2.5 nM were added to 100 μl of OPTI-MEM Medium (Thermo Fisher Scientific). The mixture was incubated at room temperature for 30 minutes prior to the addition to the coverslips.

#### Western blot analysis

Protein extracts were separated in 10% RunBlue SDS gels and transferred to PVDF membranes (Invitrogen). After blocking, membranes were incubated overnight with a peptide specific rabbit anti-IMP1 antibody (Nielsen et al., 1999) an anti-YBX1 antibody (ab12148, Abcam), an anti-GFP antibody (ab1218, Abcam) and a GAPDH antibody (FL-335, Santa Cruz) in blocking solution at 4°C before they were washed and incubated with horseradish peroxidase-conjugated anti-rabbit IgG for 1 h at room temperature. Immunoreactive proteins were detected with SuperSignal chemiluminescence reagents (Thermo Fisher Scientific) according to the manufacturer’s instructions. Blots were scanned using a C-DiGit Blot Scanner (LI-COR Biosciences).

#### Immunoprecipitation

HeLa cells were transiently transfected with pEGFP-C1 and pEGFP-IMP1 and cell pellets were collected 48 hours after transfection. Cell pellets containing 1×10^7^ cells were lysed in lysis buffer containing 20 mM Tris-HCl (pH 7.5), 1.5 mM MgCl2, 140 mM KCl, 1 mM DTT, 0.5% NP-40 supplemented with mammalian protease inhibitor cocktail (Sigma). Cell lysates were cleared by centrifugation at 8200 x*g* for 5 minutes before addition of GFP antibody (ab1218, Abcam) coupled Dynabeads^™^ Protein G (Invitrogen). Cleared cell lysates were incubated with the beads for 2 hours at 4°C with rotation. After that, samples were washed 3x with lysis buffer, split and treated with 20 ug/mL RNAse A (DNase and protease-free, Thermo Scientific) or RiboLock RNase inhibitor (Thermo Scientific) for 20 minutes at room temperature with rotation. Beads were subsequently washed and proteins were eluted directly in 2x SDS buffer.

#### Polysome fractionation analysis

Polysome analysis was performed as described (Nielsen et al., 2002). Briefly, HeLa cells (5×10^6^ cells) were lysed in 500 μl 20 mM Tris-HCl (pH 8.5), 1.5 mM MgCl_2_, 140 mM KCl, 0.5 mM DTT, 0.5% NP-40, 200 U of RNasin (Promega) per ml and 0.1 mM cycloheximide. The lysate was centrifuged at 10,000 x*g* for 10 min, and the supernatant was applied to a linear 20 to 47% sucrose gradient in 20 mM Tris-HCl (pH 8.0), 140 mM KCl, 5 mM MgCl2. Centrifugation was carried out at 200,000 *g* for 2 hours and 15 min in a Beckman SW 41 rotor. Fractions of 1 ml were collected, followed by protein precipitation with 10% TCA.

#### Electrophoretic mobility-shift analysis (EMSA)

Electrophoretic mobility-shift analysis was carried out essentially as described previously (Nielsen et al.,2004). RNA targets were *C-MYC* CRD (positions 1181-1362 in CDS), *C-MYC* 3’UTR (positions 1-226 in 3’UTR), *ACTB* exon 5, and *ACTB* 3’UTR (positions 1-233). Tag-less recombinant human IMP1 and IMP1_KH1-4mut (with the four GXXG signature loops mutated to GELG) were expressed and purified as described earlier (Nielsen et al., 2004). Recombinant human YBX1 with a C-terminal FLAG-tag was purchased from OriGene Technologies, Inc.

#### Immunofluorescence, fluorescence in situ hybridization and Structured Illumination Microscopy (SIM)

HeLa cells were seeded in glass-bottom coverslips (P35G-0.170-14-C, MatTek) and fixed 24 hours after with 3.7% formaldehyde solution in PBS, followed by a permeabilization step with 0.5% Triton X-100 in PBS. Immunofluorescence of IMP1 and YBX1 was performed using antibodies against IMP1 (E-20, Santa Cruz) and YBX1 (ab12148, Abcam). Nup153 and PABPC1 were detected using anti-Nup153 antibody (ab24700, Abcam) and anti-PABPC1 antibody (ab6125, Abcam). Coverslips were washed 3x with PBS prior to Alexa Fluor conjugated secondary antibodies (Thermo Fisher Scientific) incubation for an hour at RT. Samples were washed 3x with PBS and mounted in VECTASHIELD mounting media (RI = 1.45).

Structured Illumination Microscopy (SIM) was performed using a Zeiss ELYRA PS.1 microscope and channel correction was applied using a channel alignment file created by imaging 0.1 μm TetraSpeck Microspheres (Thermo Fisher Scientific). The image sets comprised 3 rotations and processed images were thresholded in accordance with current standards to remove diffuse background (honeycombs) caused by stray pollutants or some residual autofluorescence on the SIM reconstructed pictures before contrasting. An example is provided in Supplemental Figure 6, that also provides a comparison between conventional confocal imaging and SIM of the mRNA granules and image overlays demonstrating the measurements of the granules.

RNA FISH of *ACTB* mRNA and/or *GAPDH* mRNA was performed with human *ACTB* DesignReady Probe Set conjugated with Quasar^®^ 570 (VSMF-2002-5, LGC Biosearch) or Quasar^®^ 670 Dye (VSMF-2003-5, LGC Biosearch) or human *GAPDH* ShipReady Probe set conjugated with Quasar^®^ 570 (SMF-2026-1, LGC Biosearch) following the “Sequential Stellaris FISH and Immunofluorescence using Adherent Cells protocol” from LGC Biosearch Technologies (https://biosearchassets.blob.core.windows.net/assets/bti_custom_stellaris_immunofluorescence_seq_protocol.pdf). Briefly, immunofluroescence of IMP1 and YBX1 was performed as described above followed by a fixation step with fixation buffer (3.7% formaldehyde in PBS), Wash Buffer A (LGC Biosearch Technologies) was incubated for 5 minutes and *ACTB* probe set was hybridized in for 16 hours at 37°C with Hybridization buffer (LGC Biosearch Technologies). After hybridization, dishes were incubated with Wash Buffer A for 30 minutes at 37°C followed by addition and incubation of Wash Buffer B (LGC Biosearch Technologies) for 5 minutes. Samples were mounted and imaged using the same protocol described above.

#### OP-puromycin protein synthesis assay

HeLa cells were seeded in glass-bottom coverslips (Mattek) and Click-iT^®^ Plus OPP Alexa Fluor^®^ 488 Protein Synthesis Assay Kit was used to localize nascent polypeptides following the manufacturers protocol. Cycloheximide (50 μg/mL) was added prior to OPP to one sample as a negative control. After the click reaction, immunofluorescence detection of IMP1 and YBX1 was performed as described previously. Samples were mounted in Vectashield mounting medium and images were acquired using a Zeiss ELYRA 3.2 microscope (Structured Illumination Microscopy).

#### Photoconversion of mEos3.2-IMP1 for *in vivo* protein tracking

HeLa cells were plated in 35mm glass-bottom dishes (P35G-1.5-14-C, MatTek) and transfected with pmEos3.2-IMP1 with FuGENE 6 as described above in “Plasmid transfection” section. Photoconversion was performed using a Zeiss LSM780 microscope with a Plan-Apochromat 63x/1.4 Oil objective using a 405 laser pulse in a selected area in the cytoplasm. Cells were imaged before and after photoconversion and photoconverted mEos3.2-IMP1 was monitored every 10 seconds and followed throughout the cytoplasm.

#### Localization Microscopy

HeLa cells were stained with either IMP1 and YBX1 antibodies and Alexa Fluor 568 and Alexa Fluor 555 secondary antibodies, respectively, or with *ACTB* Quasar-570 probes (LGC Biosearch Technologies). Samples were mounted in non-hardening Vectashield mounting medium with a refraction index of 1.45 (Olivier et al.,2013). Localization Microscopy was performed on a Zeiss ELYRA PS.1 microscope using an alpha-Plan-Apochromat 100x/1.46 objective. Total internal reflection microscopy (TIRF) with a excitation wavelength of 561 nm and appropriate emission filters (BP 570-650) was used in all localization microscopy experiments and ZEN 2012 software was used to analyse and filter the data obtained in a total of 80.000 frames acquired. Frames were corrected for drift over a time scale of 36 min 13 sec (Model-based correction), and grouping was applied in the antibody stained samples in order to compile all the events that came from a single antibody. Frames corresponding to the bleaching period of the sample were discarded for the final counting.

#### Fluorescence correlation spectroscopy (FCS) and Fluorescence Cross Correlation Spectroscopy (FCCS)

FCS measurements were performed with a Zeiss LSM 780 confocal microscope. HeLa cells were transfected with pEGFP-C1 (GFP), GFP-IMP1, GFP-IMP1_KH1-4mut, GFP-YBX1 (-/+ cotransfection with YBX1 3’UTR siRNA) or PABPC1-GFP, as described above in the *Plasmid transfection* and *siRNA and plasmid cotransfection* sections and incubated for ~16 h and 48 hours respectively before measurements were conducted. Argon laser with a 488 excitation wavelength was used making sure that the count rate was linear at each particular laser power used. Transfected cells were located and FCS measurements were performed in a Zeiss LSM780 confocal microscope using a C-Apochromat 40x/1.2 W Corr M27 objective. Measurements were recorded in 10 second intervals during a total time of 60 seconds choosing arbitrary points in the cytoplasm and experimental autocorrelation curves were obtained. Intervals showing bleaching were discarded for the average. GFP-IMP1, GFP-IMP1_KH1-4mut or PABPC1-GFP were co-transfected with mCherry-YBX1, and FCCS measurements were performed following the same procedure as with FCS.

#### CLIP-seq analysis

Five public datasets were acquired by use of fastq-dump (https://trace.ncbi.nlm.nih.gov/Traces/sra/sra.cgi?view=software) using the parameters: “skip-technical”, “readids”, “read-filter pass”, “dumpbase”, “split-files”, and “clip”. Fastq files were aligned with STAR (version 2.5.2b) (Dobin et al., 2013) to the GRCh38 release 87 genome assembly as provided by ENSEMBL with corresponding annotation (ftp://ftp.ensembl.org/pub/release-87/gtf/homo_sapiens/Homo_sapiens.GRCh38.87.gtf.gz), using an overhang of 50 for all samples. Aligned reads (bam files) were imported into GenomicAlignments (Lawrence et al., 2013) in R, and single end reads were resized to the estimated fragment length (300bp) (Jothi et al., 2008). A standard peak calling algorithm could not be applied (e.g. MACS2) because of lack of a paired input sample. Hence, peaks were estimated from read pile-up (islands), where the lower boundary was estimated to be a cutoff where the mock samples proximate one island per transcript (closest number). In one case a mock was not available with the sample, and the lower boundary was estimated from median of mocks from ENCODE. All mock samples analysed required a cut-off lower than 9 reads in order to proximate one island per transcript. Crosslinked immunoprecipitation of endogenous YBX1 followed by high-throughput sequencing (CLIP-seq) in human MDA-parental breast cancer cells, YBX1 were acquired from the short read archive (SRA) with accession numbers: SRR1662159, SRR1662160, and SRR1662161 (Goodarzi et al., 2015). Enhanced CLIP-seq for IGF2BP1/IMP1 was acquired from SRA, accession numbers: SRR5112331 and SRR5112330 (Consortium,2012).

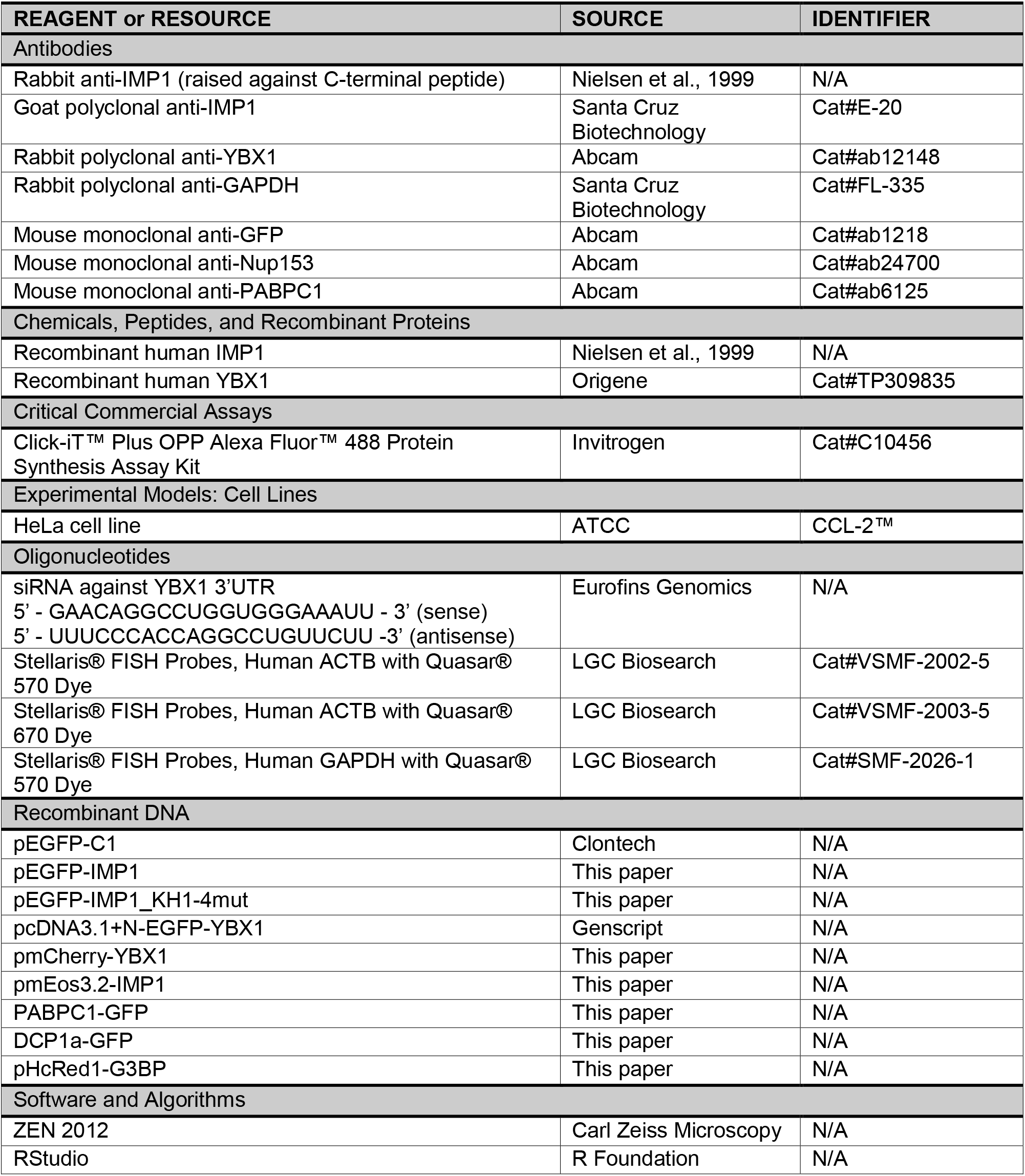
KEY RESOURCES TABLE.

