## Supplemental information for "Single mRNP analysis by super-resolution microscopy and fluorescence correlation spectroscopy reveals that small mRNP granules represent mRNA singletons"

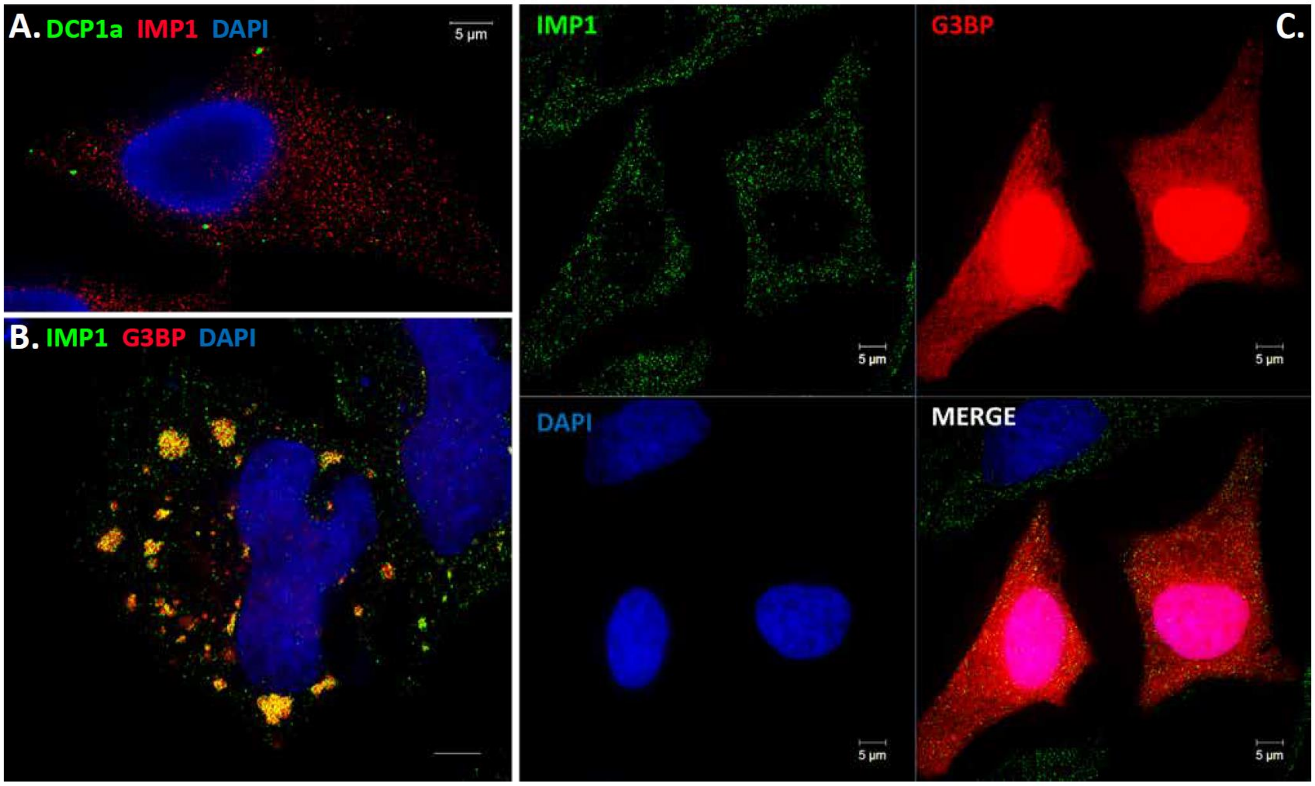


**Supplemental Figure 1**. P bodies and stress granules in combination with IMP1 **A**. P bodies depicted by DCP1a-EGFP in combination with IMP1 staining (red, Alexa Fluor 568). **B.** pHcRed-G3BP in stress granules in combination with IMP1 staining (Alexa Fluor 488, green). **C**. G3BP staining in unstressed cells showing punctuate pattern for IMP1 (green) and a diffuse pattern for G3BP (red). DAPI was employed as nuclear staining.

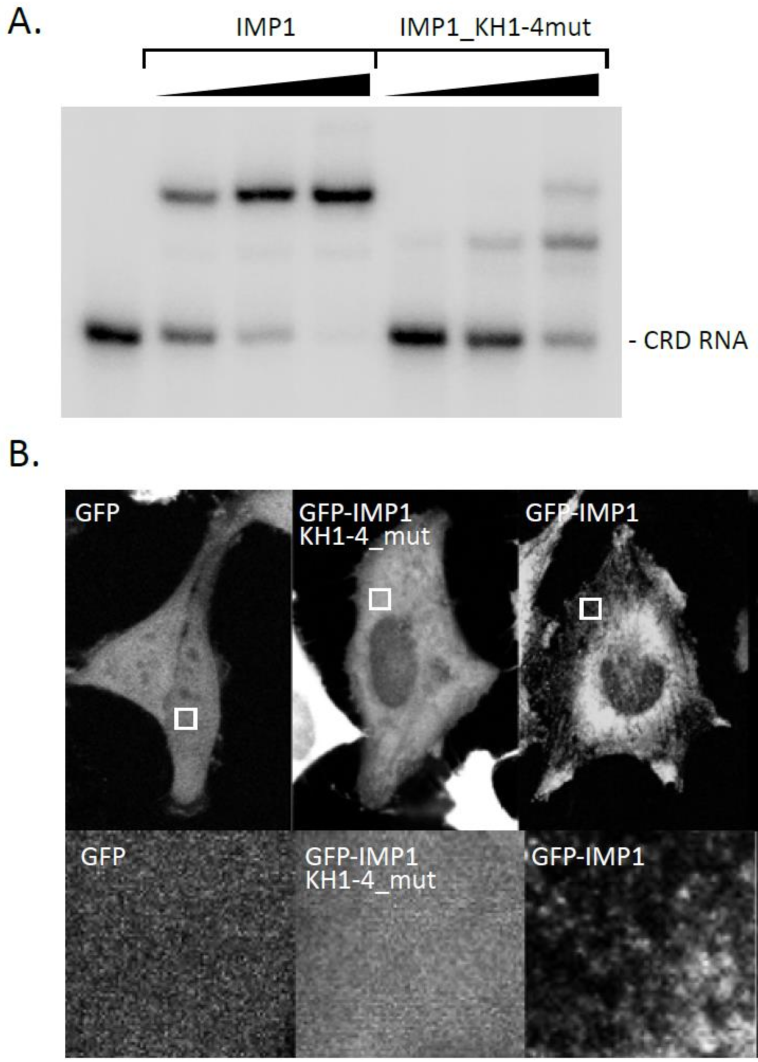


**Supplemental Figure 2. A,** Electrophoretic mobility-shift assay of IMP1 and IMP1_KH1-4mut with *C-MYC* CRD RNA probe showing that RNA-binding is impaired when IMP1 is mutated in its four KH signature loops (GXXG). There is about a 10-fold reduction in RNA-binding ability, and the dimerization of IMP1 is severely impaired. **B,** HeLa cells transfected with GFP (pEGFP-C1 vector), GFP-IMP1_KH1-4mut, GFP-IMP1 and GFP-YBX1. Upper panels show the whole cells and the lower panels show a blow up of the indicated region in the upper panels. The results show that GFP and GFP-IMP1_KH1-4mut are cytoplasmic and nuclear and are unable to form granules, whereas GFP-IMP1 and GFP-YBX1 are present in typical cytoplasmic mRNP granules.

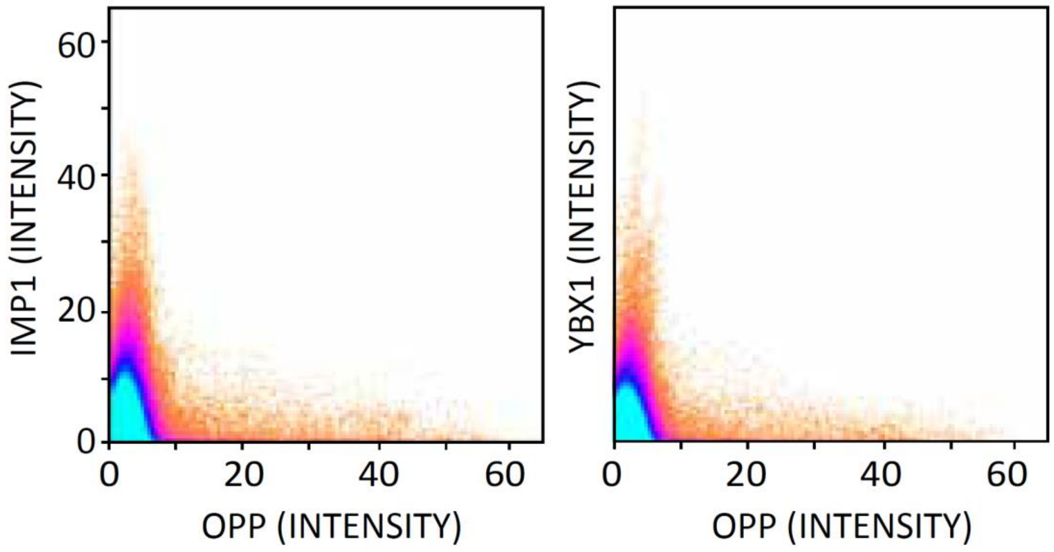


**Supplemental Figure 3**. Pixel overlaps of IMP1 vs OPP and YBX1 vs OPP.

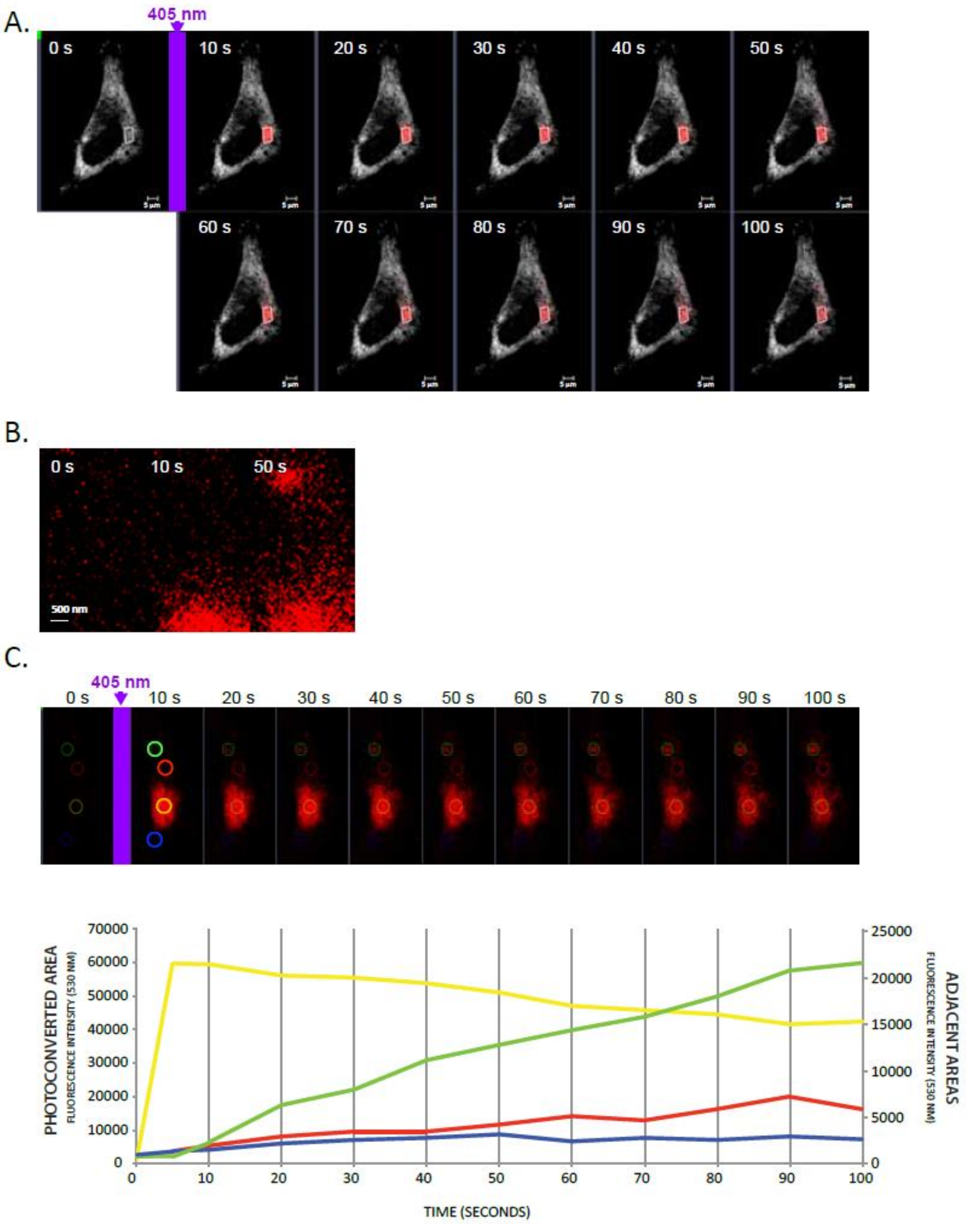


**Supplemental Figure 4. Motility of IMP1 mRNP in live cells.**

mEos-IMP1 was expressed in HeLa cells and photoconverted by a short 405 nm laser burst before migrating granules were followed by time-lapse microscopy. **A,** Time-lapse images before and after photoconversion of mEos-IMP1. Grey is mEos-IMP1 without photoconversion, whereas red is photoconverted mEos-IMP1. B. Blow up of the upper 0S, 10S and 50S frames **C,** Corresponding quantification of the fluorescence intensity of different areas (coloured circles) inside or surrounding the photoconverted area. Scale bar: 5 µm.

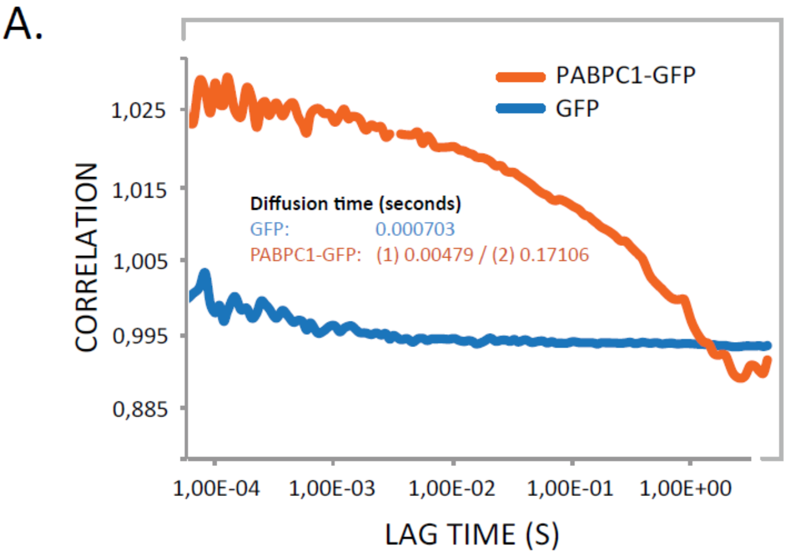


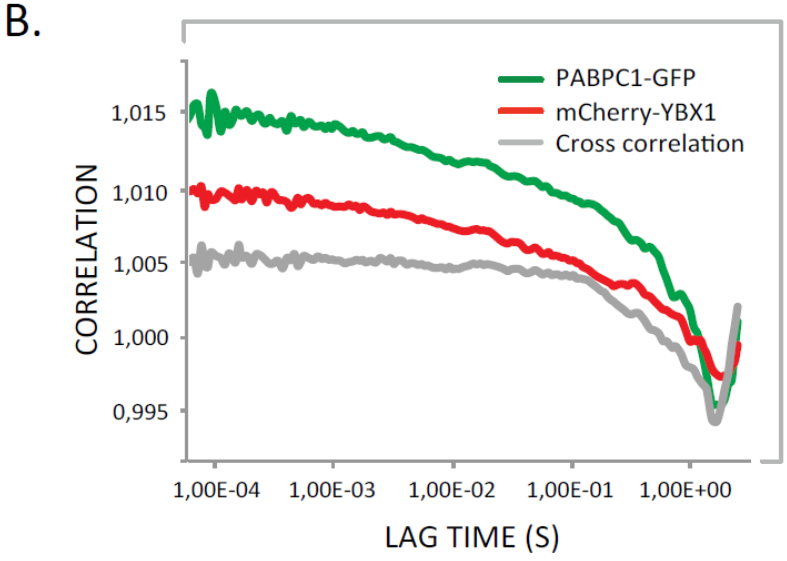


**Supplemental Figure 5. A,** GFP and PABPC1-GFP were expressed in HeLa cells and their dynamics were recorded by Fluorescence Correlation Spectroscopy (FCS). **A,** Autocorrelation curves of cytoplasmic GFP and PABPC1-GFP. Diffusion time (s) is indicated. **B,** Fluorescence Cross-Correlation curves of cells co-transfected with PABPC1-GFP and mCherry-YBX1 demonstrating interaction between the two factors *in vivo*.

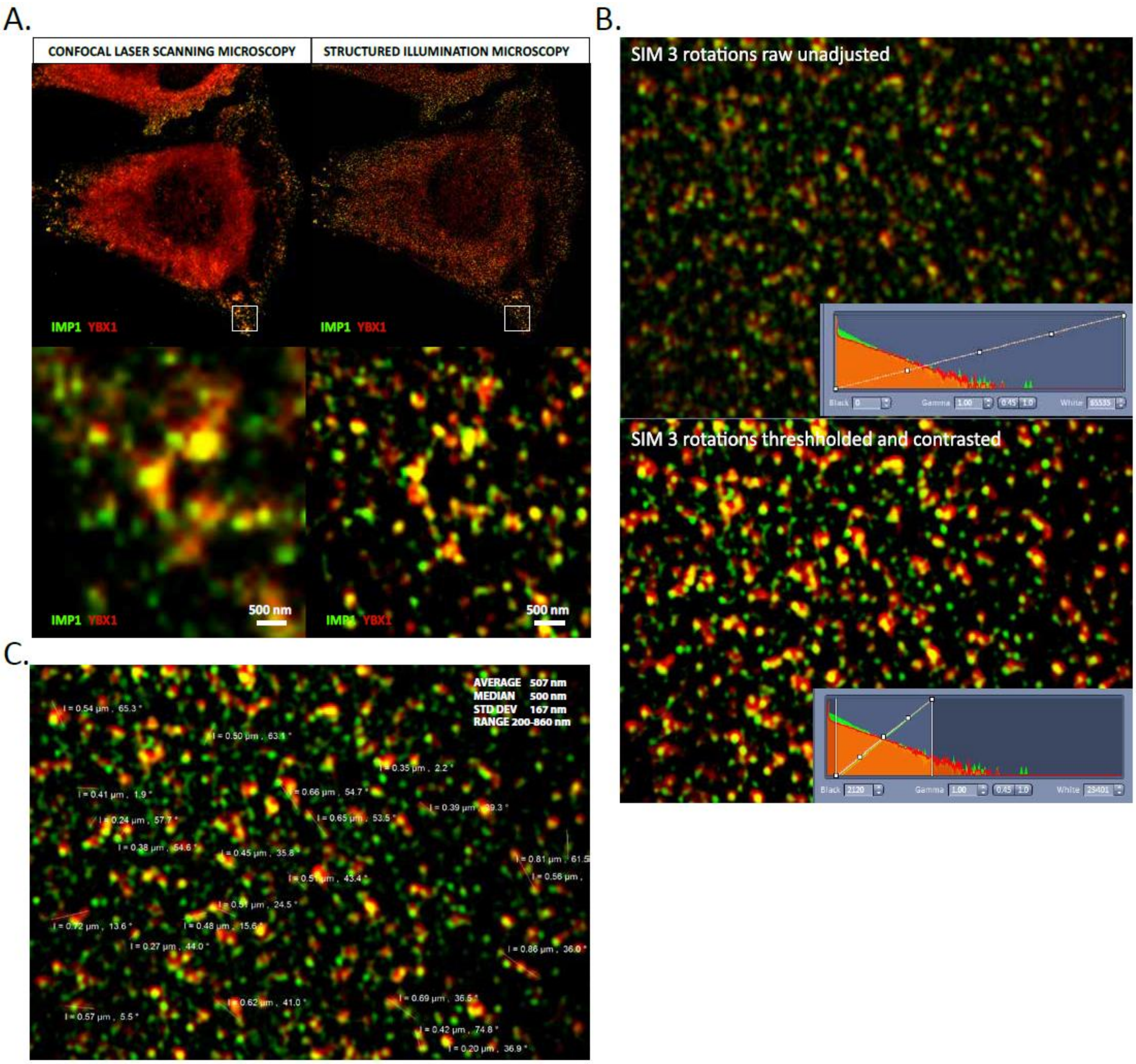


**Supplemental Figure 6. A,** Comparison between Confocal Laser Scanning Microscopy (LSM) and Structured Illumination Microscopy (SIM). HeLa cell were stained with IMP1 (green) and YBX1 (red) and imaged with LSM or SIM mode. The blow out shows how SIM is able to resolve single mRNP granules while LSM is not in most of the cases. **B.** Example of unadjusted (upper panel) and adjusted (lower panel) SIM pictures. The graph inserts show the signal intensity of each channel. Pictures were thresholded to eliminate honeycomb structures caused by caused by stray pollutants and residual autofluorescence on the SIM reconstructed pictures before contrasting. **C.** Single mRNP granule measurements.

| CLIP, eCLIP, RIP | YBX1 | | | IGF2BP1 | |
|---|---|---|---|---|---|
| CUT OFF | SRR1662159 | SRR1662160 | SRR1662161 | SRR5112330 | SRR5112331 |
| 1 | 14,46 | 11,92 | 12,82 | 31,43 | 31,19 |
| 2 | 10,19 | 8,35 | 8,65 | 30,36 | 30,13 |
| 3 | 8,31 | 7,03 | 7,08 | 16,47 | 15,52 |
| 4 | 7,12 | 6,24 | 6,17 | 15,76 | 14,59 |
| 5 | 6,25 | 5,62 | 5,52 | 9,77 | 10,10 |
| 6 | 5,58 | 5,13 | 5,01 | 9,23 | 9,38 |
| 7 | 5,04 | 4,69 | 4,60 | 6,59 | 7,56 |
| 8 | 4,62 | 4,32 | 4,24 | 6,17 | 6,97 |
| 9 | 4,28 | 4,01 | 3,94 | 4,84 | 5,98 |
| 10 | 3,99 | 3,74 | 3,68 | 4,50 | 5,52 |

**Supplemental Table 1.** “Islands per gene” analysis using different cutoffs.

| Probe | Length (nt) | Sequence |
|---|---|---|
| *C-MYC* CRD | 184 | GGACCAGAUCCCGGAGUUGGAAAACAAUGAAAAGGCCCCCAAGGUAG<br>UUAUCCUUAAAAAAGCCACAGCAUACAUCCUGUCCGUCCAAGCAGAGG<br>AGCAAAAGCUCAUUUCUGAAGAGGACUUGUUGCGGAAACGACGAGAA<br>CAGUUGAAACACAAACUUGAACAGCUACGGAACUCUUGUGCG |
| *C-MYC* 3'UTR | 226 | GGAAAAGUAAGGAAAACGAUUCCUUCUAACAGAAAUGUCCUGAGCAAU<br>CACCUAUGAACUUGUUUCAAAUGCAUGAUCAAAUGCAACCUCACAACC<br>UUGGCUGAGUCUUGAGACUGAAAGAUUUAGCCAUAAUGUAAACUGCC<br>UCAAAUUGGACUUUGGGCAUAAAAGAACUUUUUUAUGCUUACCAUCUU<br>UUUUUUUUCUUUAACAGAU UUGUAUUUAAGAAUUG |
| *ACTB* exon 5 | 183 | GGCAUGGAGUCCUGUGGCAUCCACGAAACUACCUUCAACUCCAUCAU<br>GAAGUGUGACGUGGACAUCCGCAAAGACCUGUACGCCAACACAGUGC<br>UGUCUGGCGGCACCACCAUGUACCCUGGCAUUGCCGACAGGAUGCAG<br>AAGGAGAUCACUGCCCUGGCACCCAGCACAAUGAAGAUCAAG |
| *ACTB* 3'UTR | 234 | GGCGGACUAUGACUUAGUUGCGUUACACCCUUUCUUGACAAAACCUA<br>ACUUGCGCAGAAAACAAGAUGAGAUUGGCAUGGCUUUAUUUGUUUUU<br>UUUGUUUUGUUUUGGUUUUUUUUUUUUUUUGGCUUGACUCAGGAUU<br>UAAAAACUGGAACGGUGAAGGUGACAGCAGUCGGUUGGAGCGAGCAU<br>CCCCCAAAGUUCACAAUGUGGCCGAGGACUUUGAUUGCACAUUGUU |

**Supplemental Table 2.** Sequences of the radiolabelled probes used in the Electrophoretic Mobility Shift Assay (EMSA).
